# Modulators of MAPK pathway activity during filamentous growth in *Saccharomyces cerevisiae*

**DOI:** 10.1101/2023.12.22.573138

**Authors:** Atindra N. Pujari, Paul J. Cullen

**Author notes:** Corresponding author: Paul J. Cullen Address: Department of Biological Sciences State University of New York at Buffalo Buffalo, NY 14260-1300. ANP performed experiments, wrote, and edited the paper. PJC wrote and edited the paper and obtained funding for the study.

## Abstract

Mitogen-activated protein kinase (MAPK) pathways control the response to intrinsic and extrinsic stimuli. In the budding yeast *Saccharomyces cerevisiae*, cells undergo filamentous growth, which is regulated by the fMAPK pathway. To better understand the regulation of the fMAPK pathway, a genetic screen was performed to identify spontaneous mutants with elevated activity of an fMAPK-pathway dependent growth reporter (*ste4 FUS1-HIS3*). In total, 159 mutants were isolated and analyzed by secondary screens for invasive growth by the plate-washing assay, and filament formation by microscopy. Thirty-two mutants were selected for whole-genome sequencing, which identified new alleles in genes encoding known regulators of the fMAPK pathway. These included gain-of-function alleles in *STE11,* which encodes the MAPKKK, as well as loss-of-function alleles in *KSS1,* which encodes the MAP kinase, and *RGA1,* which encodes a GTPase activating protein (GAP) for *CDC42*. New alleles in previously identified pathway modulators were also uncovered in *ALY1, AIM44, RCK2, IRA2, REG1* and in genes that regulate protein folding (*KAR2*), glycosylation (*MNN4*), and turnover (*BLM10*). C-terminal truncations in the transcription factor Ste12p were also uncovered that resulted in elevated reporter activity, presumably identifying an inhibitory domain in the C-terminus of the protein. We also show that a wide variety of filamentous growth phenotypes result from mutations in different regulators of the response. The alleles identified here expand the connections surrounding MAPK pathway regulation and reveal new features of proteins that function in the signaling cascade.

**ARTICLE SUMMARY:** Signaling pathways control the response to stimuli. In yeast, a signaling (MAPK) pathway controls a fungal behavioral response called filamentous growth. A genetic screen was performed to identify spontaneous mutants that show hyperactivity of a MAPK pathway-dependent reporter. Select mutants were analyzed by whole-genome sequencing. New alleles in known regulatory proteins were identified. A potential inhibitory domain in the C-terminus of the transcription factor Ste12p was also uncovered. Our results indicate that filamentous growth is determined by the combinatorial effects of multiple positive and negative regulatory inputs.

## INTRODUCTION

Organisms can execute a variety of responses when encountering external stimuli. The sensing and response to stimuli is facilitated by signal transduction pathways. One type of evolutionarily conserved signaling pathways are <u>m</u>itogen-<u>a</u>ctivated <u>p</u>rotein <u>k</u>inase (MAPK) pathways (Saito 2010; Bahar *et al*. 2023). MAPK pathways are found ubiquitously in eukaryotes and control the response to stress, cell differentiation, and many other responses. When mis-regulated in humans, MAPK pathways can cause diseases (Dhillon *et al*. 2007). For example, mutations that result in activation of the ERK MAPK pathway are a main cause of many types of cancers (Li *et al*. 2016). There is a strong interest in identifying proteins that regulate MAPK pathways and defining their impact on MAPK pathway activity and function.

The budding yeast *Saccharomyces cerevisiae* contains well-defined MAPK pathways (filamentous growth, mating, high osmolarity glycerol response, sporulation, and cell-wall integrity) that govern the cellular response to external stimuli (Chen and Thorner 2007; Rispail *et al*. 2009; Saito 2010). In response to limiting nutrients, like availability of carbon or nitrogen sources, the filamentous growth MAPK (fMAPK) pathway regulates a cell differentiation response called filamentous/invasive/pseudohyphal growth (Gimeno *et al*. 1992; Kumar 2021). The mating and the fMAPK pathways execute different responses yet share some components (Roberts and Fink 1994). Filamentous growth is characterized by the formation of elongated cells that undergo distal-unipolar budding, which attach and invade into surfaces in a response called invasive growth (Roberts and Fink 1994; Cullen and Sprague 2012). Filamentous growth commonly occurs in fungal pathogens (Rispail *et al*. 2009; Leach 2014) and is required for virulence some pathogens (Lo *et al*. 1997). Therefore, studying how a MAPK pathway regulates filamentous growth in *S. cerevisiae* can provide general insights into this fungal growth response.

Like other MAPK pathways, the fMAPK pathway is regulated by evolutionarily conserved factors that operate in a signaling cascade (**Fig. 1A**, **Green**). These include sensor proteins [mucin Msb2p, and sensors, Sho1p and Opy2p (Cullen *et al*. 2000; Cullen *et al*. 2004; Wu *et al*. 2006; Karunanithi and Cullen 2012; Yamamoto *et al*. 2015)], relay proteins [Rho GTPase, Cdc42; 14-3-3 proteins, Bmh1/2p; PAK, Ste20p; MAPK cascade, Ste11p, Ste7p, and Kss1p (Stevenson *et al*. 1992; Leberer *et al*. 1996; Peter *et al*. 1996; Leberer *et al*. 1997; Roberts *et al*. 1997)], and transcription factors [including Ste12p and Tec1p (Roberts and Fink 1994; Gavrias *et al*. 1996; Madhani and Fink 1997; Van der Felden *et al*. 2014)]. In addition to these positive factors, several proteins negatively regulate fMAPK pathway activity (**Fig. 1A**, **Red**). One of these proteins is Rga1p, the main GTPase-activating protein (GAP) for Cdc42p in the fMAPK pathway (Smith *et al*. 2002; Patterson *et al*. 2021). Another are the transcriptional repressors Dig1p and Dig2p (Cook *et al*. 1996; Madhani and Fink 1997; Madhani *et al*. 1997; Bardwell *et al*. 1998a; Van der Felden *et al*. 2014). The MAPK Kss1p is itself a bifunctional protein that has both positive and negative regulatory inputs (Cook *et al*. 1996; Cook *et al*. 1997).

**Figure 1.**
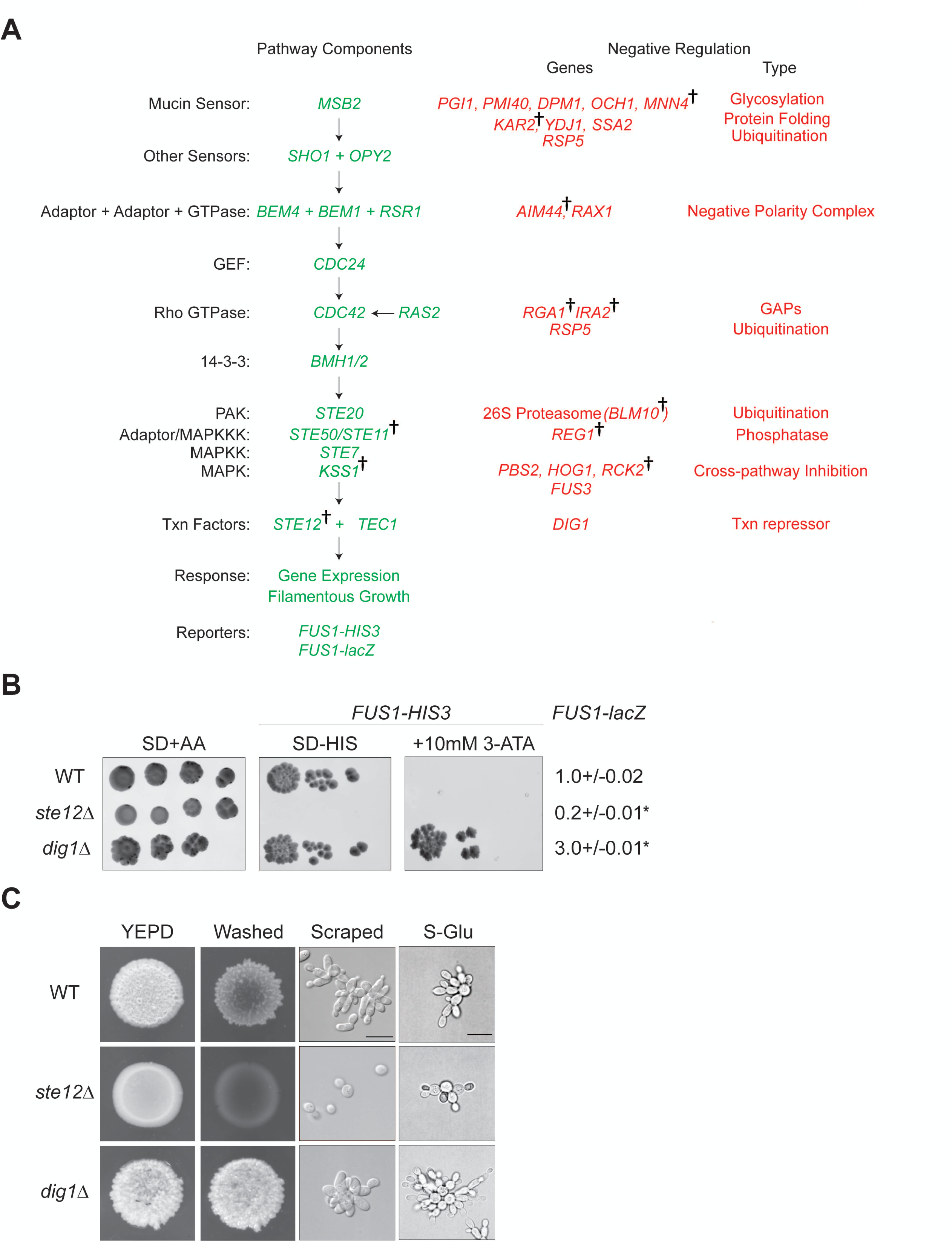
Strategy to identify modulators of the fMAPK pathway. **(A)** The fMAPK pathway. Positive components (in Green) and negative regulatory inputs (in Red) are shown. Daggers refer to genes with alleles identified in the study. **(B)** The *FUS1-lacZ* and *FUS1-HIS3* reporters were used to measure fMAPK pathway activity in WT (PC538), *ste12*Δ (PC539), and *dig1*Δ (PC3039) strains. SD+AA media denotes dilution control. SD-HIS and -HIS+10 mM 3-ATA media were used to measure fMAPK pathway activity. WT showing normal levels of fMAPK activity can grow on SD-HIS media, whereas *ste12*Δ mutant failed to grow on SD-HIS media due to downregulation of pathway activity. The *dig1*Δ mutant showing elevated activity of the pathway can grow on SD-HIS as well as on the SD-HIS+10 mM 3-ATA media. Isolates that grow on SD-HIS+10 mM 3-ATA presumably induce the fMAPK pathway. The β-galactosidase assays were performed as described. Experiments were performed in three independent replicates. Data were analyzed by one-way ANOVA followed by a Tukey’s pair-wise comparison test to generate p-values. Asterisk denotes difference compared to WT and p-value<0.05. Error bars denote standard error of the mean. **(C)** Filamentous phenotype is depicted for WT, *ste12*Δ, and *dig1*Δ strains. To compare invasive growth, strains were spotted on YEPD media, grown for 3 d at 30° C, and colonies were washed under stream of water. Colonies were photographed before and after washing. To look at cell morphology, cells were scraped and visualized using DIC at 100X magnification (scale bar, 5 microns). Cells were examined by microscopy (100X) to view the changes in cell length and budding pattern in response to glucose limitation.

Here, a genetic screen was conducted to isolate spontaneous mutants with elevated activity of an fMAPK pathway-dependent reporter, *ste4 FUS1-HIS3*, which gives a readout of fMAPK pathway activity in the absence of mating-specific genes (e.g. *STE4*). Whole-genome sequencing of select mutants identified mutations in expected genes known to impact the activity of the fMAPK pathway. The data validate previous findings that underscore key points of MAPK pathway regulation. The screen also uncovered potentially new points of regulation, including a region in the extreme C terminus of Ste12p that may interfere with the function of this homeobox transcription factor. The results furthermore reveal a diversity of phenotypes that presumably arise from combinations of mutations that impact the activity of the MAPK pathway.

## MATERIALS AND METHODS

### Yeast Strains, Media, and Growth Conditions

Strains used in the study are haploids derived from the Σ1278b strain background (Liu *et al*. 1993). Strains are listed in *Table S1* and primers in *Table S2*. Cells were grown at 30° C in YEPD, composed of yeast extract (1%), peptone (2%), and dextrose (2%), or synthetic complete (SD) media containing 0.67% yeast nitrogen base (Sigma-Aldrich, St. Louis, Missouri) and 2% glucose. Amino acids were added to the SD media as required. SD+AA denotes media containing all amino acids, SD-HIS denotes media containing all amino acids without histidine, and SD-URA denotes media containing all amino acids and lacking uracil. The His3p inhibitor 3-Amino-1,2,4-triazole (3-ATA) (Klopotowski and Wiater 1965; Struhl and Davis 1977) was added to final concentrations of 2.5 mM, 5 mM, and 10 mM as indicated. Yeast strains were manipulated using standard techniques (Rose 1990). Gene deletions were made by polymerase chain reaction (PCR) based strategies as described (Longtine *et al*. 1998). The STE4 gene deletion was made by a pop-in pop-out method (Schneider *et al*. 1995) using the integrating plasmid pSL1851 (CY2273), a pRS306-based vector. Strains containing gene deletions were checked by PCR Southern analysis.

### Genetic Screen and Characterization of Mutants

An otherwise wild-type (WT, PC538) strain lacking the *STE4* gene (PC538) was streaked onto SD+AA media for 2d at 30° C. Individual colonies were used to inoculate 5 mL of YEPD media in separate tubes. Cells were grown for 16 h at 30° C with continuous orbital shaking and were harvested (400 μL) by centrifugation. After discarding the supernatant, cell pellets were resuspended in 100 μL of autoclave-sterilized water. Cell suspension was top spread on SD-HIS+10 mM 3-ATA or SD-HIS+5 mM 3-ATA media, and plates were incubated at 30° C for 3d. Individual isolates presumably containing spontaneous mutations were collected and re-streaked on SD-HIS+10 mM 3-ATA media and grown at 30° C for three days. Strains were scraped and collected in 50% glycerol and were frozen at −80° C. Mutants were named based on the preculture number and degree of resistance to 3-ATA. Approximately, 1 × 10^8^ cells were screened for spontaneous mutations that hyperactivate the fMAPK pathway. Fifty-nine separate screens were performed to identify independent isolates.

The activity of the fMAPK pathway was measured in mutants using the *FUS1-lacZ* reporter. Mutants were further characterized by screening them for changes in invasive growth pattern by the plate-washing assay (PWA), for changes in cell morphology by differential-interference microscopy (DIC), and for changes in salt sensitivity by growth on media containing 1M KCl. Mutants were ranked from a scale of 1 (no growth on salt) to 5 (robust growth on salt) (*Table S3, Column H*). The PWA was performed as described before (Roberts and Fink 1994). Mutants were scored on a scale from 1 to 5 (1, as invasive as *ste12*Δ; 3, as invasive as WT; 5, as invasive as *dig1*Δ) (*Table S3, Column J*). For DIC images, invasive cells left behind after the PWA washes were observed at 100X using DIC filters on the Axioplan 2 fluorescence microscope (Zeiss, Jena, Germany) with a PLAN-APOCHROMAT 100×/1.4 (oil) objective (N.A. 0.17) (Zeiss, Jena, Germany). The single-cell invasive growth assay was performed as described (Cullen and Sprague 2000). WT cells (PC538) and the *ste12*Δ (PC539) and *dig1*Δ (PC3039) mutants were used as control strains.

### β-Galactosidase Assays

β-galactosidase assays were performed as described (Cullen *et al*. 2004). Cells were grown at 30° C overnight (16 h) in YEPD. Next day, 400 μL of overnight culture was centrifuged, and supernatant was discarded. The pellet was washed three times and re-suspended in 100 μL of autoclave-sterilized water. The resuspension was then used to inoculate fresh 10 mL of YEPD media. Cells were grown till mid-log phase (A_600_∼1.0), centrifuged, and pellets were stored at −80° C. Next day, pellets were resuspended in 100 μL Z-buffer [44.32 mL H_2_O with 5 mL phosphate buffer (0.6 M Na_2_HPO_4_ + 0.4 M Na_2_HPO_4_), 0.5 mL 1 M KCl, 50 μL 1M MgSO_4_, 135 μL β-mercaptoethanol]. Once resuspended, 2 μL of 5% Sarkosyl (S) and 2 μL of toluene were added, and the pellets were incubated at 37° C for 30 min with open caps to allow for evaporation of toluene. Then, Z+S buffer containing ortho-Nitrophenyl-β-galactosidase was added to the pellets. After color change was observed, reactions were stopped by adding 250 μL of 1M Na_2_CO_3_, and reaction time was recorded. Reaction mixture was centrifuged (13,000 rpm for 3 min) to remove cell extract, and 200 μL of supernatant was used to determine A_420_. (1000 X A_420_)/(A_600_ X time) was used to calculate miller units. Miller Units demonstrated in graphs are averages of three or more independent experiments along with the standard error of the means.

### Genomic DNA Extraction, Sequencing Analysis, and Identification of Variants

For generating chromosomal DNA for sequencing analysis, the Gentra Puregene Yeast Bacteria DNA extraction kit was used (Qiagen, Hilden, Germany). Cells were grown for 16 h in 5 ml cultures and prepared according to manufacturer’s protocols. Sequencing and variant calling were performed at the Genomics & Bioinformatics shared resource of the Fred Hutchinson Cancer Center in Seattle Washington. Genomic DNA was quantified using Life Technologies’ Invitrogen Qubit 2.0 Fluorometer (Thermo Fisher, Waltham, MA) and fragmented using a Covaris LE220 ultrasonicator (Covaris, Woodburn, MA) targeting 400 bp. Sequencing libraries were prepared using 100 ng fragmented DNA with the KAPA HyperPrep Library Prep Kit (Roche, Indianapolis, IN) and NEXTFLEX UDI barcodes (PerkinElmer, Waltham, MA). Library quantification was performed using Life Technologies’ Invitrogen Qubit® 2.0 Fluorometer and size distribution validated using an Agilent 4200 TapeStation (Agilent Technologies, Santa Clara, CA). Individual libraries were pooled 30-plex at equimolar concentrations and sequenced on one lane of an Illumina HiSeq 2500 (Illumina, Inc, San Diego, CA) employing a paired-end, 50-base read length sequencing configuration. This yielded 3M to 5.5M (average 4.5M) read pairs covering the ∼12Mb *S. cerevisiae* genome per sample.

Read processing and germline variant calling followed GATK best practice workflow (https://gatk.broadinstitute.org/hc/en-us/articles/360035535932-Germline-short-variantdiscovery-SNPs-Indels-). Briefly, sequencing reads were mapped to yeast genome assembly sacCerS1278b (strain Stanford S1278b, https://www.yeastgenome.org/strain/sigma1278b) using BWA 0.7.17 (Li and Durbin 2009). Resulting alignments, in BAM format, were further processed using GATK 4.1.4.1 (Mckenna *et al*. 2010; Van der Auwera *et al*. 2013) to generate analysis-ready alignments with proper read group information and duplicated reads marked (MarkDuplicates). Next, per-sample variant calling was conducted using the GATK HaplotypeCaller in GVCF mode followed by joint calling of all samples using CombineGVCFs and GenotypeGVCFs. Initial calls were filtered using VariantFiltration with the following parameters: QUAL < 20.00 MQ < 30.00, GQ < 5.0 and DP < 6.0. Filtered variants were annotated using snpEff (Cingolani *et al*. 2012).

Variant sites among all 34 samples (includes a pooled sample) with respect to the Σ1278b reference genome were identified (*Table S4A)*. To identify differences between wild type and the mutant lines, results were filtered to 556 variants where at least one mutant line had a genotype different from both the reference genome and the wild type. The resulting variant calls (in VCF format) were compiled into a single table providing an overview of detected variants (*Table S4B*, one row per variant). A custom database was constructed for running snpEff, a tool to annotate variants, exported in html format. snpEff takes GATK generated VCF file and generates a new VCF file with updated INFO field (snpEff.annotation/filtered.ann.vcf). The format is described in http://snpeff.sourceforge.net/SnpEff_manual.html#input. The relevant information was consolidated to snpEff.annotation/filtered.variants.annotation.xlsx. Each tab “legend” shows a description of each field. Intragenic variants identified in each mutant were compiled into a single sheet to show the genes, alternate alleles, amino acid change, and mutants that contain the alternate alleles (*Table S4C*).

### Phospho-MAP Kinase Analysis

Cells were inoculated in YEPD or SD+AA media (5 mL) and grown for 16 h at 30° C with continuous shaking. Approximately 750 µl of cells from the saturated culture grown for 16 h were inoculated into 10 mL SD+AA media, and cells were grown until the culture reached mid-log phase (A_600_ ∼ 1.0). Five mL of cells were harvested from mid-log phase cultures for immunoblot analysis. Cells were disrupted, and proteins were enriched by tri-chloroacetic acid precipitation as previously described (Basu *et al*. 2016). Protein samples were analyzed by sodium dodecyl sulfate polyacrylamide gel electrophoresis (SDS-PAGE) analysis and were transferred from gels to a nitrocellulose membrane (Cat#10600003, Amersham Protran Premium 0.45 μm NC; GE Healthcare Life sciences). Membranes were probed with rabbit polyclonal p44/42 antibodies (Cell Signaling Technology, Danvers, MA; Cat #4370) diluted 1:10,000 in 5% bovine serum albumin (BSA) to detect P∼MAP kinases, P∼Fus3p and P∼Kss1p. Monoclonal mouse anti-Pgk1p antibodies (Life Technologies; Camarillo, CA; Cat #459250) were used at 1:10,000 dilution as a control for total protein levels. Secondary anti-mouse IgG-HRP (Cat#1706516; Bio-Rad, Inc.) and goat anti-rabbit IgG-HRP (Cat#115-035-003; Jackson ImmnunoResearch Laboratories) were used to detect the primary antibodies. The nitrocellulose membrane was blocked with 5% non-fat dried milk for Pgk1p anti-body or 5% BSA for the p44/42 antibody for 1 h prior to antibody incubation. Primary antibody incubations were performed at 4° C for 16 h, and secondary antibody incubations were performed at 22° C for 1 h. Immunoblots were visualized by Gel Doc XR Imaging System (Bio-Rad, Inc.), after addition of Chemiluminescent HRP substrate for chemiluminescent Westerns (Radiance Plus Substrate, Azure Biosystems). Raw images are included for the blots shown in (**Fig. S5**; A, membrane probed with p44/42 antibodies; and B, membrane probed with Pgk1p antibodies).

Densitometric analysis was performed with Image Lab Software (Bio-Rad). The same exposures in the linear range were used for measuring band intensities. Background subtraction was performed according to the guidelines provided by the manufacturer. Band intensities of phospho-proteins were normalized against total protein levels based on Pgk1p band intensity.

### Statistical Analysis

All statistical tests were performed with Minitab, LLC. 2021. Retrieved from https://www.minitab.com. Data were analyzed by one-way ANOVA test followed by a Tukey’s or Dunnett’s (control group, WT) multiple comparison tests to generate p-values.

## RESULTS and DISCUSSION

### Strategy to identify modulators of the fMAPK pathway

The fMAPK pathway is composed of proteins that promote (**Fig. 1A, Green)** or inhibit (**Fig. 1A, Red)** MAPK pathway activity. In cells lacking an intact mating pathway (*ste4*), transcriptional reporters *FUS1-HIS3* and *FUS1-lacZ* (**Fig. 1B**, both integrated into the genome) provide a readout of fMAPK pathway activity (Cullen *et al*. 2004). With respect to the *FUS1-HIS3* reporter, WT cells (PC538) grow on SD-HIS media, whereas cells lacking an intact fMAPK pathway [e.g. the *ste12*Δ (PC539) mutant] do not. By comparison, cells lacking negative regulators of the pathway, such as the *dig1*Δ (PC3039) mutant grow on media lacking histidine and containing 3-ATA, which is a competitive inhibitor of His3p enzyme (**Fig. 1B**). Therefore, mutants that grow on SD-HIS+3-ATA media presumably show elevated fMAPK pathway activity. Likewise, WT cells show basal *FUS1-lacZ* reporter expression by β-galactosidase assays (**Fig. 1B**). The activity of this reporter is also dependent on the transcription factor Ste12p and modulated by the transcriptional repressor Dig1p (**Fig. 1B**).

To observe aspects of filamentous growth (invasion, morphology, and budding pattern), WT cells along with the *ste12*Δ and *dig1*Δ mutants were spotted onto YEPD media and grown for 3 d at 30° C. Colonies were washed off the plate to reveal invasive scars **(Fig. 1C**, Washed) by the plate-washing assay (PWA) (Roberts and Fink 1994). The *ste12*Δ mutant showed less agar invasion than WT cells, whereas the *dig1*Δ mutant showed elevated invasive growth **(Fig. 1C**, Washed). Microscopic examination showed that WT cells were elongated and remained attached to each other, whereas *ste12*Δ mutant cells were round and separated from each other. The *dig1*Δ mutant cells showed increased cell elongation **(Fig. 1C**, Scraped). By the single cell assay (Cullen and Sprague 2000), WT cells showed distal-pole budding, whereas the *ste12*Δ mutant showed reduced distal budding. The *dig1*Δ mutant cells showed hyper cell polarization (**Fig. 1C**). Therefore, the filamentous growth phenotypes controlled by the fMAPK pathway parallel transcriptional reporter activity.

### Genetic screen and characterization of mutants

A genetic screen was performed to identify genes that when mutated stimulate the activity of the fMAPK pathway reporter (*ste4 FUS1-HIS3*). Expected mutations include loss-of-function mutations in genes encoding negative regulators and gain-of-function mutations in genes encoding positive regulators. Spontaneous mutants were isolated that were capable of growth on SD-HIS+10 mM 3-ATA media. In total, 159 mutants were identified in 59 separate batches and were phenotypically characterized (*Table S3*). Separate batches were used to eliminate duplicate mutations. Of the 159 mutants, 144 grew on SD-HIS+10 mM 3-ATA media and 15 grew on SD-HIS+5 mM 3-ATA media (*Table S3, Column D*). To eliminate potential contaminants, isolates were streaked on SD-URA media. No URA+ isolates were uncovered (*Table S3, Column K*), indicating that the mutants were derived presumably from the *ura3*-parent strain (*Fig. S1A*).

Examining the mutants on the same media showed that some mutants grew better on SD-HIS+10 mM 3-ATA than others (**Fig. 2**, inset). To quantitate MAPK pathway levels, the activity of the *FUS1-lacZ* reporter was determined by β-galactosidase assays. Most mutants showed elevated levels of reporter activity (**Fig. 2**, bar graph). Mutants that did not show elevated *FUS1-lacZ* activity may contain mutations in the *ATR1* gene which causes 3-ATA resistance or mutations in the *FUS1-HIS3* promoter and were not considered further.

**Figure 2.**
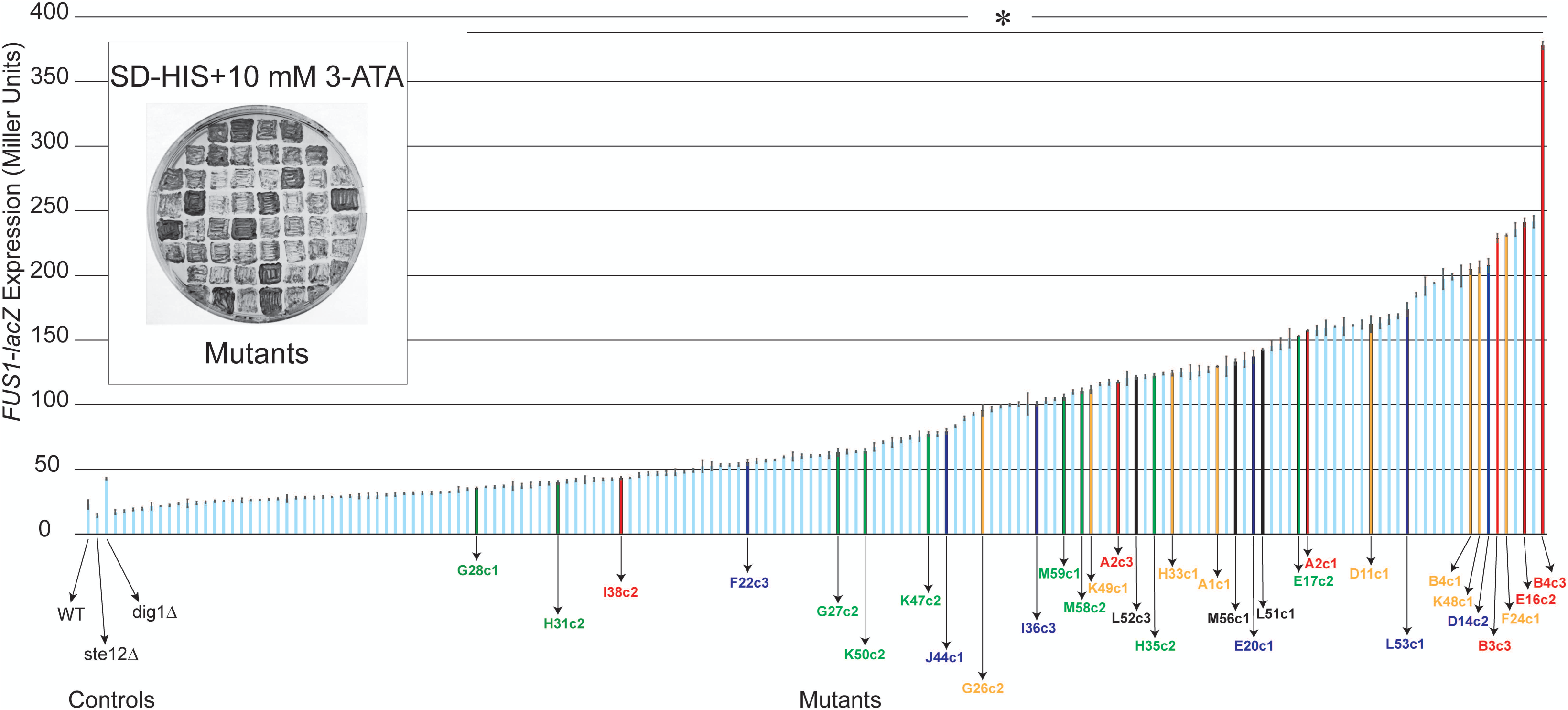
Genetic screen identifies mutants that hyperactivate the fMAPK pathway. Inset, mutants that show hyperactivity of the *FUS1-HIS3* reporter; a subset of fifty-five mutants that grow on SD-HIS+10 mM 3-ATA media are shown. Validation of fMAPK pathway activity by the *FUS1-lacZ* reporter. WT (PC538), *ste12*Δ (PC539), and *dig1*Δ (PC3039) strains were used as controls. Experiments were performed in three independent replicates. Data were analyzed by one-way ANOVA test followed by a Dunnett’s multiple comparison test to generate p-values. Asterisk denotes significant difference compared to WT and p-values of <0.05. Error bars denote standard error of the mean. Differences in *lacZ* levels between experiments (e.g. Fig. 1B) may result from different growth conditions. The thirty-two indicated mutants were chosen for sequencing analysis. Colors represent mutations in the indicated genes: red, *KSS1*; green, *RGA1*; blue, *STE11*; and orange, *STE12*.

Mutants were further screened by additional tests. To eliminate known mutations in the high osmolarity glycerol response (HOG) pathway that are known to cause activation of the fMAPK pathway [specifically *pbs2* and *hog1* (O’rourke and herskowitz 1998)], mutants were screened for salt sensitivity by growth on media containing 1M KCl. Mutants showed varying levels of salt sensitivity (*Fig. S1B*; *Table S3*, Column H). Mutants showing a growth defect on salt were not selected for further analysis.

Mutants were also examined by the PWA to identify changes in invasive growth (**Fig. 3**, the complete dataset can be found in *Fig. S2A-D*; *Table S3*, Column J). Invasive cells were also examined by microscopy (**Fig. 3**, right panels; the complete dataset can be found in *File S1*). A subset of mutants showed growth defects on YEPD media (*Table S3*, Column G), which might arise due to strong hyperactivity of the fMAPK pathway. To capture a broad collection of genotypes, a phenotypically diverse collection of mutants was selected for whole-genome sequencing analysis (*Table S3*, Column O).

**Figure 3.**
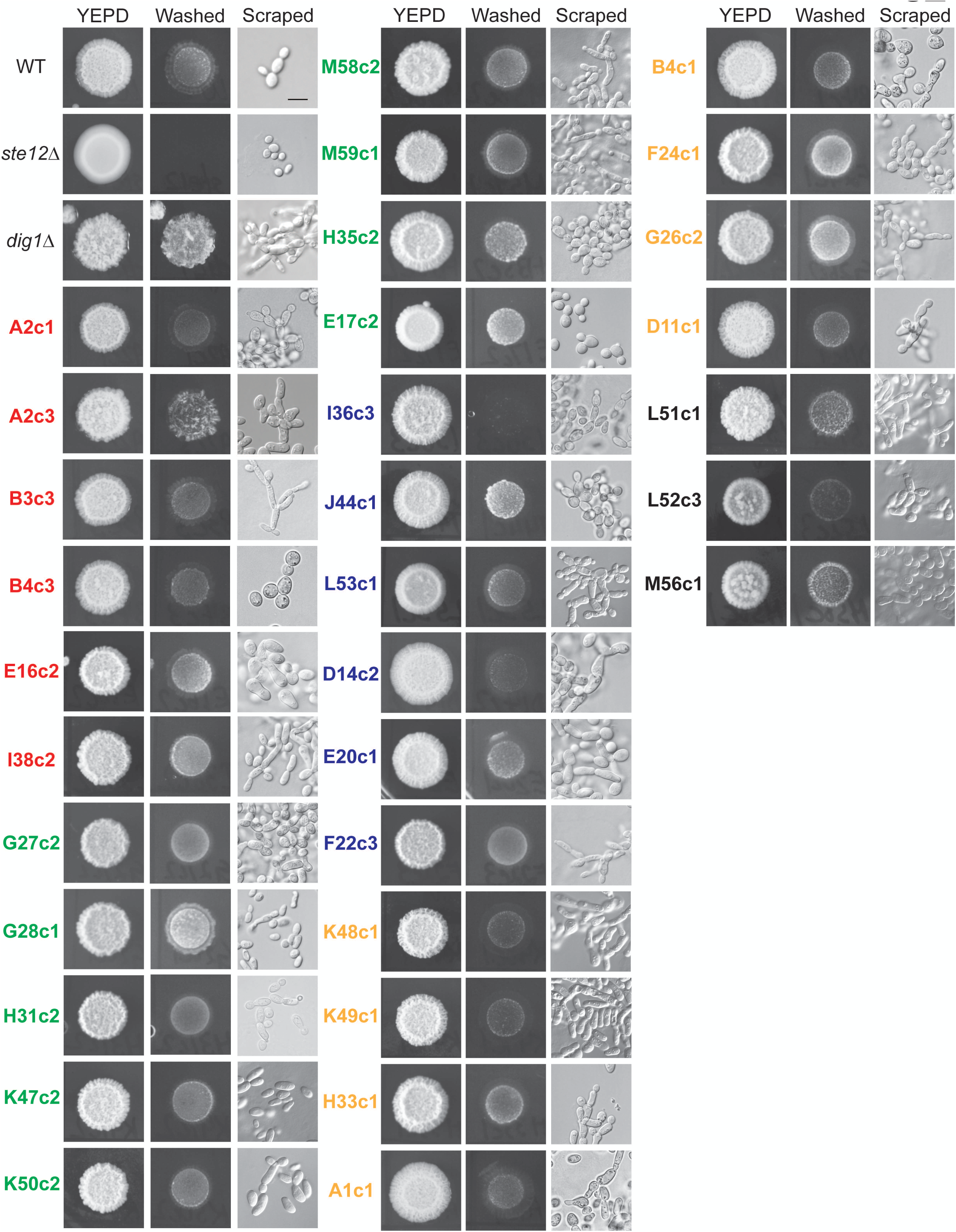
Secondary screens of mutants for invasive growth and cell morphology phenotypes. Plate-washing assay was used to assess invasive growth phenotypes. Mutants were spotted in 5 μL volume on YEPD media and incubated at 30° C for 3 d. Plates were washed under a stream of water to reveal invasive scars. Photographs were taken before and after washing. The full dataset is shown in *Fig. S2A-D*. Invaded cells were scraped off of washed plates by a toothpick and observed at 100X magnification using DIC filter. Bar, 10 microns. A typical example of WT (PC538), *ste12*Δ (PC539), and *dig1*Δ (PC3039) from the experiment are shown as controls. Colors represent mutations in the indicated genes: red, *KSS1*; green, *RGA1*; blue, *STE11*; and orange, *STE12*.

### Whole-genome sequencing and bioinformatics analysis to identify variants

For sequencing analysis, DNA was harvested from thirty-two mutants and the WT control strain by chromosomal preparation. DNA sequencing was performed with an Illumina HiSeq 2500 machine, which yielded an average of 4.5M read pair coverage over the yeast genome. The total number of variants identified compared to the reference genome was 2,131 (*Table S4A*). The variant rate was one variant every 4,943 base pairs and included single nucleotide polymorphisms, additions, and deletions. Overall, 196 missense, 21 nonsense, and 461 silent mutations were recovered. The missense to silent ratio was 0.4252.

To better define the relevant changes in the mutants, the variations in each mutant were compared to somatic variants that differed between the wild-type parental strain and the reference genome, Σ1278b. As a result, 557 mutations were identified across the mutants sequenced (*Table S4B*). The mutations mapped to ORFs as well as intergenic regions upstream and downstream of the ORFs (*Table S4B*). We focused on the mutations present in exons (*Table S4C*), which revealed a set of mutations expected to impact the activity of the fMAPK pathway. The mutations identified here validate the utility of the screen by recovery of expected mutations surrounding the reporters. Many of the alleles (the term mutation and allele are used interchangeably) remain uncharacterized and could contribute to some of the phenotypic variation observed here. In addition, structural variation in genes (repeat regions) may not have been detected by short sequencing reads and might also contribute to phenotypic variation. These features could be examined in future analysis.

### Alleles uncovered by analysis of DNA sequencing data

The screen identified mutations in genes previously known to regulate the fMAPK pathway (*Table 1*). These included mutations in 4 genes encoding known regulators of the pathway Ste11p, Rga1p, Kss1p, and Ste12p (**Fig. 1**, Daggers). Multiple alleles in each of the genes were uncovered, and each gene is discussed in detail below.

**Table 1.**
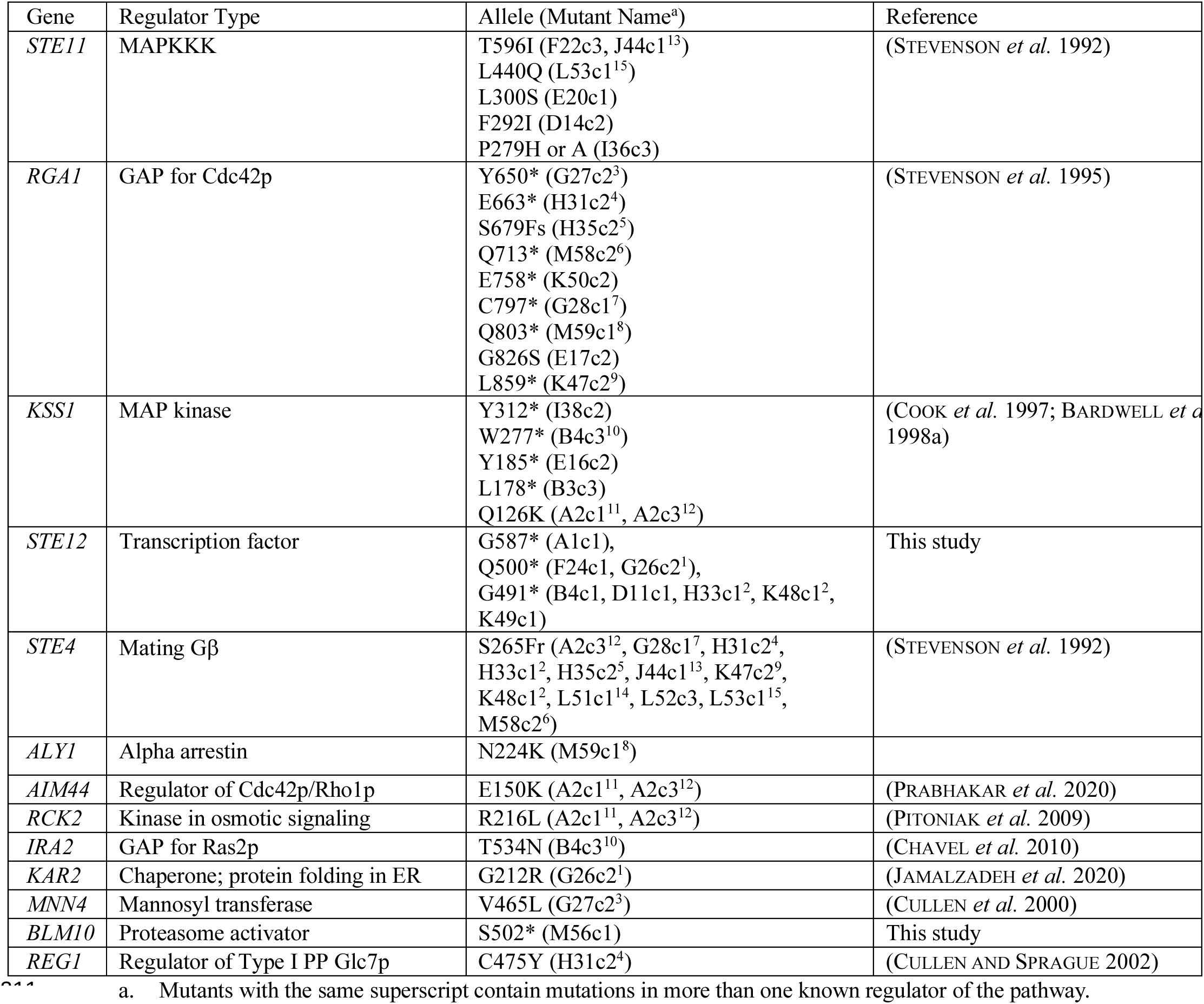
Mutations identified in genes regulating the fMAPK pathway.

#### STE11

Six mutants contained mutations in the *STE11* gene (*Table 1*). Two missense mutations led to changes at the P279H and P279A position. A previous study uncovered an allele that resulted in the P279S change, called *STE11-1* (Stevenson et al. 1992), which hyperactivated the MAPKKK to similar levels as we saw here. Another gain-of-function mutation was uncovered that resulted in the change T596I, which was previously uncovered as the strong hyperactive *STE11-4* allele (Stevenson *et al*. 1992). Three new mutations were also identified in the central region of the protein upstream of the kinase domain that would be predicted to change the following residues, L440Q, L300S, and F292I. Although not tested, these changes might also be expected to impact the activity of the Ste11p protein.

#### RGA1

Nine mutants contained mutations in the gene encoding Rga1p, the main GAP for Cdc42p that functions in the fMAPK pathway Rga1p (*Table 1*). Eight of the mutations had frameshifts (S679Fr) or premature stop codons (Y650*, E663*, Q713*, E758*, C797*, Q803*, and L859*). These mutations would be expected to result in loss-of-function of the Rga1p protein, as the GAP domain in the C-terminus of the protein would be expected to be removed. Loss of GAP activity is expected to result in elevated Cdc42p activity, resulting in higher levels of Cdc42p-dependent fMAPK pathway activity (Stevenson *et al*. 1995). One missense mutation was uncovered resulting in the amino acid substitution G826S, which changed a conserved glycine residue found in other Rho GAPs including Bem2p, Bem3p, and Lrg1p (Stevenson *et al*. 1995). Therefore, this change might also be expected to compromise GAP activity of the Rga1p protein.

#### KSS1

Six mutants had mutations in the gene encoding the MAPK Kss1p (*Table 1*). Four were premature stop codons (Y312*, W277*, Y185*, and L178*). Two mutants contained the same missense mutation that led to the change Q126K. Loss-of-function alleles in the MAPK Kss1p may be expected to be uncovered because Kss1p has both positive and negative regulatory functions (Cook *et al*. 1997; Bardwell *et al*. 1998a). As shown below, loss of Kss1p function is known to result in elevated phosphorylation of a related MAP kinase, Fus3p, which would be expected to induce fMAPK pathway reporter activity.

#### STE12

Eight mutants contained mutations in the gene encoding the transcription factor Ste12p. These mutations all contained premature stop codons that would be expected to result in C-terminal truncations of the protein. Five mutants contained G491*, two mutants contained Q500*, and one mutant contained G587*. Loss of Ste12p would be expected to result in reduced reporter activity (**Fig. 1B**). Therefore, loss of the C-terminal domain of Ste12p is presumably inhibitory, leading to elevated fMAPK pathway activity.

#### Other Genes

Twelve mutants contained mutations in *STE4*, which might explain their hyperactive phenotypes. *STE4+* revertant mutants were presumably recovered by repair of the *STE4* gene. Overall, of the 32 mutants sequenced, 31 mutants had mutations in one of the above-mentioned genes (*STE11*, *RGA1*, *KSS1*, *STE12*, and *STE4*), which could explain the increased reporter activity found in these strains. The remaining mutant contained a premature stop codon, S502*, in *BLM10*, which encodes a non-essential gene that functions as the activator of the 19S proteasome (Dange *et al*. 2011). The proteasome is required for turnover of Cdc42p (Gonzalez and Cullen 2022) and Ste20p (Gonzalez *et al*. 2023), which are positive regulators of the fMAPK pathway. Therefore, cells lacking the ability to degrade these proteins might show elevated activity of the fMAPK pathway.

Although multiple alleles of *STE11*, *RGA1*, *KSS1*, and *STE12* were uncovered in the screen, we did not find any examples of a mutant containing mutations in more than one of these genes. This may be because multiple mutations in genes that hyperactive the fMAPK pathway cause growth defects or otherwise show genetic interactions which prevent their detection by this approach. Comparing mutants that contained different alleles of above-mentioned genes revealed unexpected variation in reporter activity (**Fig. 2**, compare orange to orange, etc) and in filamentous growth phenotype (**Fig. 3**, compare mutants of the same color). We speculate that this may be due to phenotypic differences in the alleles for a given gene. Alternatively, phenotypic differences may arise due to mutations in other genes and intergenic regions. Variation in phenotype caused by alleles of the same gene may also be due to epigenetic factors that lead to stochastic differences within strains (Kaern *et al*. 2005; Zhang *et al*. 2013).

To identify the genetic basis of this variation, we looked for other mutations that might be expected to impact reporter activity. Several candidate genes were uncovered and are discussed here. A mutation in *ALY1* resulting in N224K amino acid change was uncovered. When overexpressed, the gene encoding the alpha arrestin, *ALY1* (O’donnell et al. 2013), dampens the fMAPK pathway activity due to mislocalization of the sensor protein Msb2p (Adhikari *et al*. 2015a). A mutation in *AIM44/GPS1* resulting in E150K amino acid was also uncovered. Loss of *AIM44*, a member of the negative polarity complex (Meitinger *et al*. 2014), is known to cause hyperactivity of the fMAPK pathway (Prabhakar *et al*. 2020). A mutation in *RCK2* resulting in R216L amino acid change was also identified. *RCK2* encodes a calmodulin-like kinase and a target of Hog1p, (Bilsland-Marchesan *et al*. 2000; Teige *et al*. 2001), which dampens fMAPK pathway activity when overexpressed (Pitoniak et al 2008). We also uncovered mutations in genes involved in protein folding [*KAR2*, G212R; (Rose *et al*. 1989)], and glycosylation [*MNN4*, V465L; (Odani *et al*. 1996)] in the endoplasmic reticulum. Defects in protein folding and glycosylation of Msb2p are known to hyperactivate the fMAPK pathway (Cullen *et al*. 2000; Pitoniak *et al*. 2009; Adhikari *et al*. 2015b; Jamalzadeh *et al*. 2020). A mutation in *IRA2* resulting in T534N was uncovered. The *IRA2* gene encodes a GAP for the GTPase Ras2p (Halme *et al*. 2004). Loss of *IRA2* would be expected to increase Ras2p activity, which itself is a positive regulator of the fMAPK pathway (Mosch *et al*. 1996). A mutation in *REG1* resulting in C475Y was uncovered. *REG1* encodes a regulator of type I protein phosphatase Glc7p (Sanz *et al*. 2000). The *REG1-GLC7* phosphatase complex is also known to negatively regulate glucose repressible genes. Glc7p is known to regulate filamentous growth (Cullen and Sprague 2002) and the mutations identified here may also impact fMAPK pathway activity.

Several other alleles were also identified and tested for a role on fMAPK pathway reporter activity. Gene deletions were constructed (*nam7*Δ*, aly1*Δ*, bnr1*Δ*, esc1*Δ*, yke2*Δ*, pac10*Δ*, ssb1*Δ*, ena1*Δ*, tor1*Δ, and *snf5*Δ) and tested for altered levels of *FUS1-HIS3* reporter activity. None of the mutants tested showed hyperactivity of the pathway based on this test (*Fig. S3*). Apart from the above-mentioned mutations that were recovered in non-essential genes, several mutations were identified in essential genes (*Table S4C,* Column J). The roles that essential genes play in filamentous growth and fMAPK pathway activity are being explored in a separate study (Pujari *et al*. in preparation).

### Examination of select mutants by P∼Kss1p analysis

To validate and extend these findings, the activity of the fMAPK pathway was measured in select mutants by phospho-immunoblot analysis. The level of phosphorylated Kss1p (P∼Kss1p) provides a readout of fMAPK pathway activity. The same antibodies (anti-p44/42) that recognize P∼Kss1p also recognize P∼Fus3p, which mainly regulates the mating pathway. Because several genes identified here are shared among MAPK pathways, the alleles identified in this study could have an effect on several MAPK pathways. The WT (PC538), *ste11*Δ (PC611) and *STE11-4* (PC580) strains were used as controls to observe basal, defective, and hyper P∼Kss1p/P∼Fus3p levels, respectively. When comparing mutants to WT and the *ste11*Δ control, all the mutants showed hyper phosphorylation of either Kss1p or Fus3p or both MAP kinases suggesting that mutations in these mutants lead to hyperactive MAPKs. The mutant F22c3 containing T596I, which is the same substitution in Ste11-4p (Stevenson *et al*. 1992), caused similar hyperactivation of P∼Kss1p and P∼Fus3p as seen in *STE11-4* control. Like the Ste11-1p variant (P279S) identified previously (Stevenson *et al*. 1992), the mutant I36c3 (P279H/A) showed hyper phosphorylation of Kss1p and Fus3p. Different alleles leading to specific amino acid substitutions in Ste11p and present in different mutants showed unique patterns of P∼Kss1p and P∼ Fus3p levels (E20c1, L300S; I36c3, P279H/A; L53c1, L440Q; D14c2, F292I; and F22c3, T596I). Like the *STE11* alleles, the *RGA1* allele leading to C-terminal truncation of the protein (E758*) present in the K50c2 mutant showed hyper phosphorylation of Kss1p and Fus3p. Interestingly, all three mutations in the *KSS1* gene leading to premature stop codons, L178* and Y185*, present in B3c3 and E16c2, respectively, and Q126K, present in A2c1, show low levels of P∼Kss1p and high levels of P∼Fus3p. The mutations leading to truncated Kss1p might cause reduced protein levels, which would explain loss of P∼Kss1p in those mutants. However, it is unclear how the missense mutation Q126K causes a defect in P∼Kss1p levels. Because Kss1p is known to have a negative regulatory function in the fMAPK pathway (Cook *et al*. 1997; Bardwell *et al*. 1998a), loss of Kss1p might lead to Fus3p*-* dependent activation of *FUS1* reporter. In support of this idea, loss of Kss1p did not result in hyper invasive growth in these mutants by the PWA (**Fig. 3**). Two mutants, H33c1 and K48c1, containing the same mutation that leads to premature stop codon at the G491 position in Ste12p, also caused similar hyper phosphorylation of Kss1p and Fus3p. This may indicate that the C-terminal domain of Ste12p is not required for fMAPK pathway activation.

### Comparison of fMAPK pathway activity to invasive growth and cell morphology

Many of the mutants identified in the study showed higher *FUS1-lacZ* reporter activity than WT (**Fig. 2**, asterisk denotes significant difference). However, *FUS1-lacZ* reporter activity did not always correlate with invasive growth (**Fig. 3**) or P∼Kss1p levels (**Fig. 4**). Similarly, the polarized morphology of filamentous cells did not always correlate with invasive growth. Therefore, we directly compared the effects of known hyperactive mutations on invasive growth, cell morphology, and *FUS1-lacZ* reporter activity (**Fig. 5**). All the hyperactive mutants tested showed upregulation of *FUS1-lacZ* reporter. The *rga1*Δ mutant showed a modest increase in reporter activity compared to the *STE11-4*, *pmi40-101,* and the *dig1*Δ mutants (reporter activity in ascending order, *ste12*Δ < WT < *rga1*Δ < *pmi40-101* < *dig1*Δ < *STE11-4*). By comparison, the *dig1*Δ mutant was more invasive than other hyperactive mutants and the WT and *ste12*Δ controls (invasion in ascending order, *ste12*Δ < *pmi40-101* < WT = *STE11-4* < *rga1*Δ < *dig1*Δ). When comparing cell morphology, the *dig1*Δ mutant showed the most cell polarization compared to other mutants and controls (cell polarization in ascending order, *pmi40-101* < *ste12*Δ < WT < *STE11-4* = *rga1*Δ < *dig1*Δ). Thus, the output phenotypes of filamentous growth did not match each other or the activity of the fMAPK pathway. This may be because different regulators of the fMAPK pathway might control different aspects of filamentous growth. The fact that different hyperactive alleles lead to different output phenotypes makes it difficult to assign genotypic information based on phenotype.

**Figure 4.**
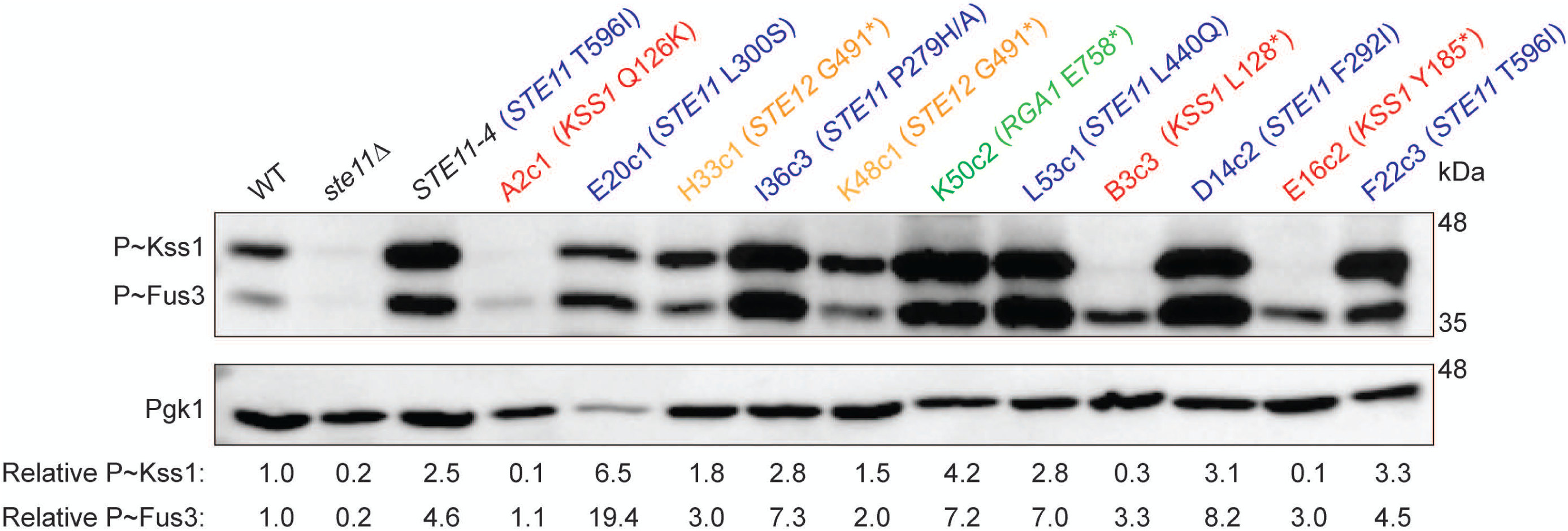
Analysis of the levels of phosphorylated Kss1p in selected mutants. Immunoblots of membranes probed with p44/42 antibodies to visualize P∼Kss1p and P∼Fus3p levels, and Pgk1p antibodies as a control for total protein levels. WT (PC538), and *ste11*Δ (PC611) and *STE11-4* (PC580) strains were used as standards. Band intensity was quantified using ImageLab. Phopho-band intensities were normalized to Pgk1p bands for each lane, and the WT phospho-protein levels were set to one to compare P∼MAP kinases in different mutants with respect to WT. Colors represent mutations in the indicated genes: red, *KSS1*; green, *RGA1*; blue, *STE11*; and orange, *STE12*.

**Figure 5.**
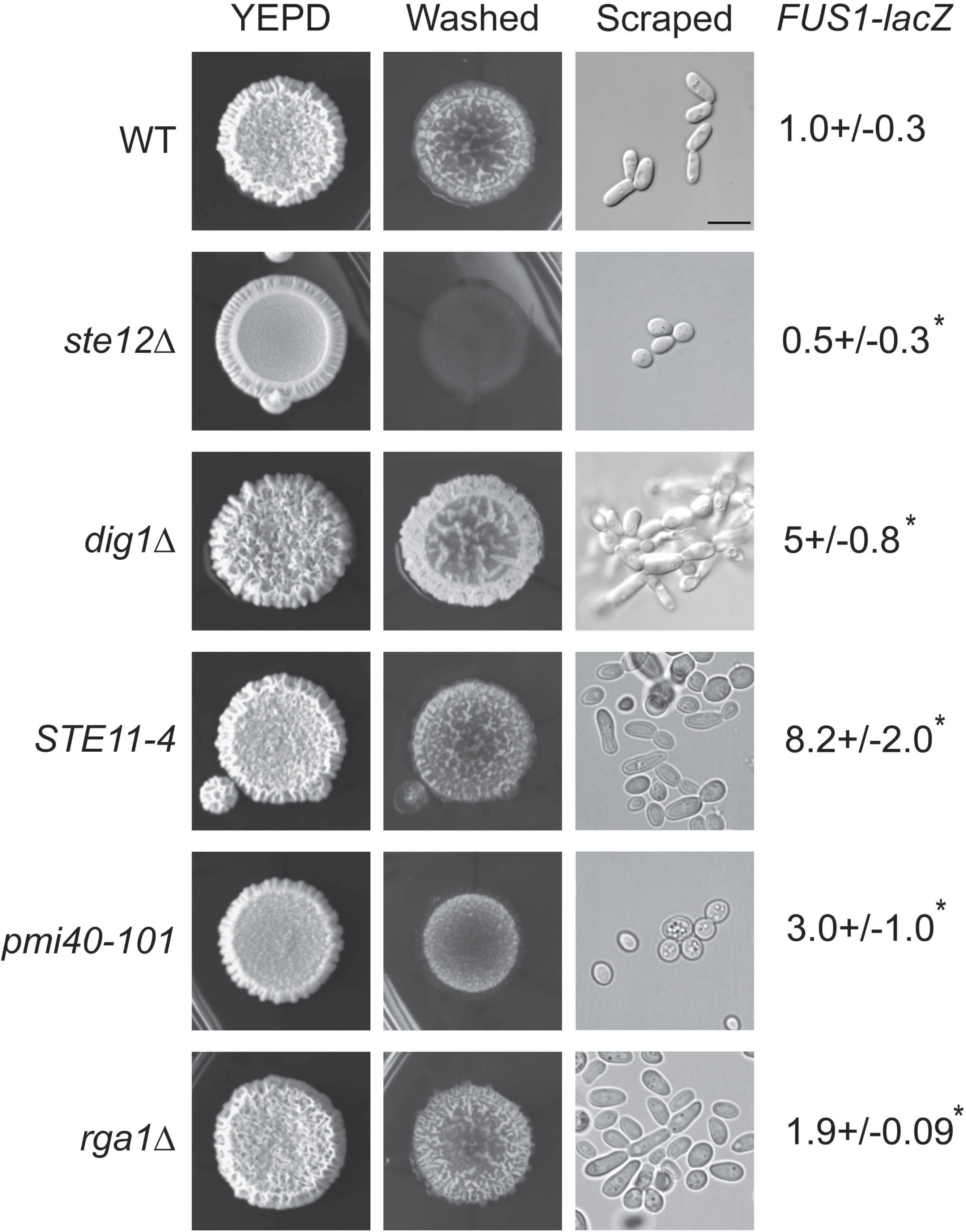
Comparison of phenotypes of mutants that hyperactivate the fMAPK pathway. Strains were spotted in 5 μL volume on YEPD plates, grown for 3 d at 30° C, and washed under stream of water to reveal invasive scars. Pictures were taken before and after washing. Invasive cells were scraped and observed at 100X magnification using DIC filter. For *FUS1-lacZ* assays, strains were grown for 16 h in YEPD media, sub-cultured into YEPD, and grown till mid-log phase (A_600_ ∼ 1.0). Mid-log phase cells were collected to prepare lysates for β-galactosidase assays. WT (PC538) values were compared with values for mutant defective for fMAPK activity (*ste12*Δ, PC539) or hyperactive for fMAPK activity (*dig1*Δ, PC3039; *ste11-4*, PC580; *rga1*Δ, PC3391; and *pmi40-101*, PC5014). Experiments were performed in three independent replicates. Data were analyzed by one-way ANOVA test followed by a Tukey’s pair-wise comparison test to generate p-values. Asterisk denotes differences compared to WT, p-value<0.05. Error bars denote standard error of the mean. Differences in *lacZ* levels between experiments (e.g. Fig. 1B) may result from different growth conditions.

Here, a genetic screen was performed to identify mutants that show elevated activity of a MAPK pathway-dependent reporter. Whole-genome sequencing of a subset of mutants identified new alleles of well-established regulators of the MAPK pathway. These included gain-of-function alleles that induce constitutive activation of Ste11p, several of which had been identified previously (Stevenson *et al*. 1992). Likewise, loss-of-function mutations in *RGA1*, encoding the GAP for Cdc42p, are similar to those previously described (Stevenson *et al*. 1995). Although not previously identified, loss-of-function alleles predicted to truncate the MAP kinase Kss1p were also uncovered. These alleles might be expected to cause elevated activity of the fMAPK pathway, because Kss1p has an inhibitory role (Cook *et al*. 1997; Bardwell *et al*. 1998a). Unexpectedly, mutations predicted to truncate the C-terminal domain of Ste12p (from G491, E500, and G587) were also uncovered. We infer that the C terminus of Ste12p may function to inhibit the activity of this factor or alter its interaction with interacting partners. Multiple alleles of the above-mentioned genes were identified. By comparison, alleles in other genes (*ALY1*, *AIM44, RCK2, BLM10, KAR2, MNN4, IRA2,* and *REG1*) were uncovered at a low frequency that have been previously shown to impact the activity of the fMAPK pathway. At least some of these inputs presumably come from other pathways, such as the Ras2-cAMP-PKA pathway. Ras2p is known to regulate the fMAPK pathway (Mosch *et al*. 1996), and we identified mutations in the gene encoding for a Ras2p GAP, Ira2p, here. Furthermore, defects in protein glycosylation are also known to stimulate the activity of the fMAPK pathway (Cullen *et al*. 2000).

We show here that mutations resulting in loss of C-terminal domain of Ste12p are functional and show higher levels of MAPK pathway activity than the wild-type protein. Ste12p is a member of the homeodomain family of transcription factors (Rispail and Di PIETRO 2010). Ste12p homologs exist in many fungal species and have been characterized in plant and animal pathogens (Rispail and Di PIETRO 2010; Wong SAK HOI and dumas 2010; Fan *et al*. 2021; Purohit and Gajjar 2022). In plant pathogens, *Colletotrichum lindemuthianum and Botrytis cinerea*, the extreme C terminus of Ste12p is modified by alternative splicing (Wong SAK HOI *et al*. 2007; Schamber *et al*. 2010). Depending on the spliced products formed, different combinations of zinc-finger domains are produced at the end of the transcription factor, resulting in activating or inhibitory forms of the Ste12p proteins. The yeast Ste12p lacks Zn finger domains. Ste12p interacts with other proteins that lead to changes in gene expression. One group of proteins are the transcriptional repressors Dig1p and Dig2p (Tedford *et al*. 1997; Bardwell *et al*. 1998a; Bardwell *et al*. 1998b; Zheng *et al*. 2010). Moreover, Ste12p can impact the organization of gene expression in the nucleus (Randise-Hinchliff *et al*. 2016), which occurs through a mechanism that involves the post-translational modification of Dig2p (Randise-Hinchliff *et al*. 2016). In addition, Ste12p recognizes a specific consensus sequence on the DNA to regulate mating specific genes, and a different motif when in complex with Tec1p to control invasive growth (Madhani and Fink 1997; Heise *et al*. 2010; Dorrity *et al*. 2018; Zhou *et al*. 2020). Ste12p and Tec1p also regulate the cyclin-dependent kinase Cdk8p, controlling RNAPII-C-terminal domain to maintain genome integrity (Nelson *et al*. 2003; Aristizabal *et al*. 2015). It will be interesting to determine if the C terminus of Ste12p interacts with any of these factors and how this might impact fMAPK pathway signaling.

Gain-of-function mutations in *STE11* were also identified in this study. Mutations in *STE11* were compared to mutations in RAF, which encodes the MAPKKK that regulates the ERK MAP kinase pathway in humans (Wellbrock *et al*. 2004). Mutations in RAF are associated with constitutive activation of the protein and cancer (Turski *et al*. 2016). Alignment of the proteins revealed a region in the kinase domain of Ste11p (VKI**T**DFGISKK) that shows homology to the same region of B-RAF (VKIG**DFG**GLATV). In Ste11p, T596I identified here and previously [*STE11-4*, (Stevenson *et al*. 1992)] leads to constitutive activation of the kinase domain of the protein. Similarly, in B-RAF, changes to the adjacent amino acids D, F, and G have been linked to cancer. The valine at position 600 in B-RAF, which is not conserved in Ste11p, is the most changed residue found in cancer patients (Turski *et al*. 2016). A second site in Ste11p (<u>PS</u>EF) is conserved with (<u>PS</u>KS) in B-Raf, which corresponds to the previously identified *STE11-1* allele [P279S, (Stevenson *et al*. 1992)] and the two alleles identified in this study (P279H/A). The other changes in Ste11p identified here (L440Q, L300S, and F292I) did not show homology to regions in the RAF protein. Therefore, at least two regions in MAPKKKs are conserved between yeast and humans, which when changed results in presumptive elevated kinase activity and phenotypic effects.

Some expected mutations were not identified by the screen. Internal deletions in the N-terminal inhibitory domain of Msb2p, which leads to elevated fMAPK pathway activity (Cullen *et al*. 2004), were not recovered. A version of transmembrane sensor Sho1p, Sho1p^P120L^, which hyperactivates fMAPK pathway (Vadaie *et al*. 2008), was also not uncovered by the screen. It is unclear why mutations in some genes were recovered multiple times (*STE11, STE12, RGA1* and *KSS1*), while other mutations were under-represented (e.g. in *ALY1, IRA2*), and still other mutations were not recovered (e.g. in *TEC1, STE20, STE7*). Perhaps some regions of the genome are more susceptible to spontaneous mutations than others. Alternatively, growth defects of some mutants may mask the pathway activity. These could include alleles in essential genes, like those controlling protein glycosylation. Other mutants (e.g. *dig1,* which was not found) could induce enhanced cell adhesion and polarized cell morphologies that may impact colony morphology and were ignored. However, many mutations were uncovered in intergenic regions and upstream and downstream of the ORFs that were not explored here. Moreover, mutations that result in elevated MAPK pathway activity can lead to mutations in other parts of the genome, as retrotransposon mobility can be induced by the fMAPK pathway (Company *et al*. 1988; Conte *et al*. 1998).

## Data Availability

All strains and mutants described in the study are available upon request to. The authors affirm that all the data supporting the conclusions of the study can be found in the article, figures, and tables. Supplemental data is available at FigShare. Raw DNA sequences have been deposited on a publicly available resource.

## ACKNOWLEDGEMENTS

Thanks to Matthew Fitzgibbon, Feinan Wu, and other members of the Fred Hutchinson Genome Resource Center for DNA sequencing and variant analysis. Thanks to laboratory members for suggestions, and Ankita Priyadarshini for reading the manuscript.

## ABBREVIATIONS

3-ATA, 3-Amino-1,2,4-triazole; BSA, bovine serum albumin; DIC, differential interference contrast; ER, endoplasmic reticulum; HOG, high osmolarity glycerol; MAPK, mitogen-activated protein kinase; PWA, plate-washing assay; PCR, polymerase chain reaction; Rho, RAS homology; S, Sarkosyl; SD, synthetic complete; SDS-PAGE, sodium dodecyl sulfate polyacrylamide gel electrophoresis; YEPD, yeast extract, peptone, and dextrose

